# Novel mutations associated with autosomal dominant congenital cataract are identified in Chinese families

**DOI:** 10.1101/367516

**Authors:** Zhenyu Wang, Chen Huang, Yanxiu Sun, Huibin Lv, Mingzhou Zhang, Xuemin Li

## Abstract

**Purpose:** As the leading cause of the impairment of vision of children, congenital cataract is considered as a hereditary disease, especially autosomal dominant congenital cataract (ADCC). The purpose of this study is to identify the genetic defect of six Chinese families with ADCC.

**Subjects and Methods:** Six Chinese families with ADCC were recruited in the study. (103 members in total, 96 members alive, 27 patients in total) Genomic DNA samples extracting from probands’ peripheral blood cells were captured the mutations using a specific eye disease enrichment panel with next generation sequencing. After initial pathogenicity prediction, sites with specific pathogenicity were screened for further validation. Sanger sequencing was conducted in the other individuals in the families and other 100 normal controls. Mutations definitely related with ADCC will then be analyzed by bioinformatics analysis. The pathogenic effect of the amino acid changes and structural and functional changes of the proteins were finally analyzed by bioinformatics analysis.

**Results:** Seven mutations in six candidate genes associated with ADCC of six families were detected (MYH9 c.4150G>C, CRYBA4 c.169T>C, RPGRRIP1 c.2669G>A, WFS1 c.1235T>C, CRYBA4 c.26C>T, EPHA2 c.2663+1G>A, and PAX6 c.11–2A>G). All the seven mutations were only detected on affected individuals in the families. Among them there are three novel mutations (MYH9 c.4150G>C, CRYBA4 c.169T>C, RPGRRIP1 c.2669G>A) and four that have been reported (WFS1 c.1235T>C, CRYBA4 c.26C>T, EPHA2 c.2663+1G>A, and PAX6 c.11–2A>G). RPGRIP1 (c.2669G>A) mutation and CRYBA4 (c.26C>T) mutation are predicted to be benign according to bioinformatics analysis while the other five mutations (EPHA2, PAX6, MYH9, CRYBA4 c.169T>C, WFS1) are thought to be pathogenic.

**Conclusion:** We report two novel heterozygous mutations (MYH9 c.4150G>C and CRYBA4 c.169T>C) in six Chinese families supporting their vital roles in causing ADCC.

## Introduction

During the initial developmental period of vision, lens opacity mainly caused by congenital cataract may lead to form deprivation amblyopia [1]. According to the studies, about one third of blindness of infants is caused by congenital cataracts [2,3]. The incidence of congenital cataract is 6.31/100,000[4] in industrialized countries while the figure of that in developing countries is assumed to be much higher [5,6].

As is known as the leading cause of children’s impairment of vision globally, the hereditary modes of congenital cataract include autosomal dominant, autosomal recessive and X-linked hereditary mode. Among all the modes, autosomal dominant congenital cataract (ADCC) is the most common [4], which is also known that variable clinical features are presented in different families [7,8]. Until now, at least 23 genes have been associated with ADCC. These genes are mainly involved in the formation of the lens. They are classified as crystallin genes (CRYA, CRYB, and CRYG), lens specific connexin genes (Cx43, Cx46, and Cx50), major intrinsic protein gene (MIP) or aquaporine, cytoskeletal structural protein genes, paired-like homeodomain transcription factor 3 (PITX3), avian musculoaponeurotic fibrosarcoma (MAF), heat shock transcription factor 4 (HSF4), beaded filament structural protein 2(BFSP2) and non-muscle myosin heavy chain IIA, NMMHC-IIA (MYH9)[7,9,10]. Besides, Ephrin receptor subfamily (EPHA1, EPHA2), RPGR interacting protein 1 (RPGRIP1) and paired box 6 (PAX6) also play a vital role in pathogenesis of cataract.

However, mutations in the same gene may lead to different phenotype [7,8]. Even for individuals in the same family, they may have ADCC with different clinical features. As we found, variable clinical features occurs in the patients in the same family as well as different families. In this study, we conducted targeted gene capture using a hereditary-eye-disease-enriching panel and next generation sequencing to identify the mutations of six Chinese families with ADCC. According to this study, two novel mutations in MYH9 (c.4150G>C) and CRYBA4 (c.169T>C) were detected. The mutations were not detected in the unaffected individuals in the Family 1, Family 4 and 100 normal controls. It means that these mutations may be pathogenesis and responsible for ADCC.

## Subjects and Methods

### Recruitment of patients and clinical evaluation

Owing to seeking the gene therapy for the next generation of the probands (III-1 in Family 1; IV-7 in Family 2; IV-1 in Family 3; III-4 in Family 4; III-2 in Family 5; III-2 in Family 6), six Chinese families with 103 members in total (96 members alive) were recruited from the Peking University Third Hospital (**Figure 1**). There are 27 patients in total who have ADCC in the family (4 patients from Family 1; 6 patients from Family 2; 4 patients from Family 3; 7 patients from Family 4; 3 patients from Family 5; 3 patients from Family 6). Detailed family and medical histories and a series of results of ophthalmic examinations were obtained from the family members, including visual acuity, slit lamp examination and fundus examination with dilated pupils. All participating individuals provided informed consents consistent with the tenets of the Declaration of Helsink. Peking University Third Hospital Medical Ethics Committee approved all procedures used in this study.

**Figure 1.**
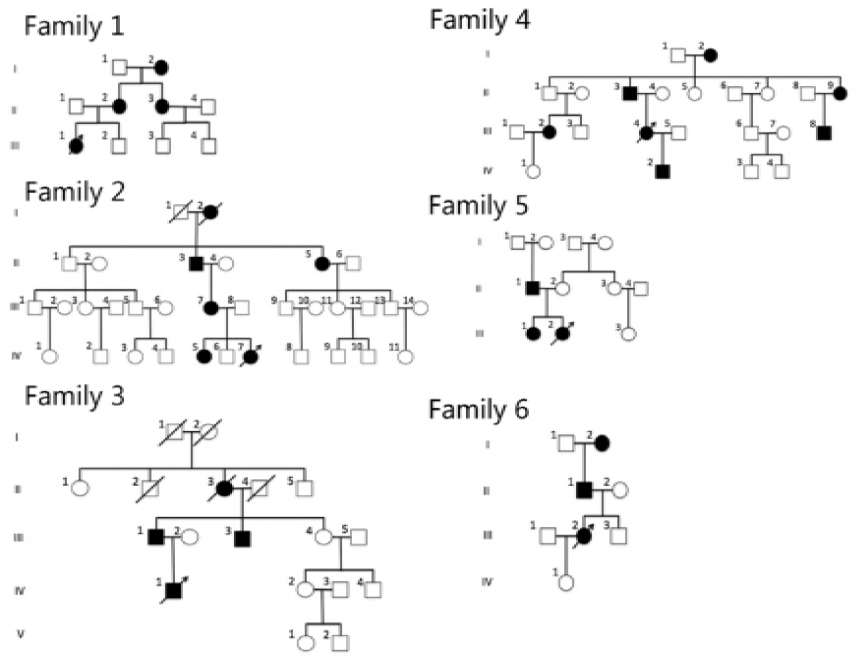
Pedigree of six families with autosomal dominant congenital cataract (ADCC).
Squares and circles indicate males and females, respectively. Blackened symbols denote
affected status.

### Genomic DNA Extraction

Two milliliters of venous blood was collected from the participating family members and was stored in BD Vacutainers (BD, San Jose, CA) which contains EDTA to prevent coagulation. The genomic DNA was extracted from the white blood cells using DNA Extraction kit (TIANGEN, Beijing, China), and was quantified with Nanodrop 2000 (Thermal Fisher Scientific, DE).

### Mutation Screening

After extracting DNA from the white blood cells of the proband (III-1 in Family 1; IV-7 in Family 2; IV-1 in Family 3; III-4 in Family 4; III-2 in Family 5; III-2 in Family 6), a specific eye disease enrichment panel was used to capture the mutation of gene sample (MyGenostics GenCap Enrichment technologies Inc., Beijing, China). A minimum of 3 microgram DNA was used for the indexed Illumina libraries according to manufacturer’s protocol (MyGenostics, Inc., Beijing, China). The target genes in the enriched libraries were captured by using MyGenostics Targeted Genes Capture protocol and then were sequenced on an llumina NextSeq 500 sequencer (Illumina, San Diego, CA, USA) for paired-end reads of 150 bp.

A total of 663 disease genes in the panel have relationship with hereditary eye diseases. Among these genes, there are 135 genes (**Table 1**) associated with cataract (57 genes are associated with congenital cataract and others are associated with hereditary eye diseases with opacified lens).

**Table 1.** 135 cataract-associated genes that are in the panel

|  |  |  |  |  |  |
| --- | --- | --- | --- | --- | --- |
| ABHD12 | ADAMTSL4 | AGK | AGPS | AIPL1 | ALDH18A1 |
| ATOH7 | BCOR | BEST1 | BFSP1 | BFSP2 | CAPN5 |
| CAV1 | CEP290 | CHMP4B | CNBP | COL11A1 | COL11A2 |
| COL2A1 | COL7A1 | COL9A1 | COL9A2 | CRB1 | CRX |
| CRYAA | CRYAB | CRYBA1 | CRYBA2 | CRYBA4 | CRYBB1 |
| CRYBB2 | CRYBB3 | CRYGA | CRYGB | CRYGC | CRYGD |
| CRYGS | CTDP1 | CYP27A1 | CYP51A1 | DMPK | DNA2 |
| EPG5 | EPHA1 | EPHA2 | ERCC2 | ERCC5 | ERCC6 |
| EZR | FAM126A | FTL | FYCO1 | GALK1 | GALT |
| GCNT2 | GDF6 | GFER | GJA3 | GJA8 | GNPAT |
| GSTM1 | GSTT1 | GUCY2D | HFE | HMX1 | HSF4 |
| IARS2 | IDO1 | IFNGR1 | IMPDH1 | ITM2B | JAM3 |
| LCA5 | LCT | LIM2 | LMX1B | LRAT | MAF |
| MIP | MVK | MYH9 | NAT2 | NECTIN3 | NEUROD1 |
| NF2 | NHS | NMNAT1 | NOD2 | OAT | OCRL |
| OPA3 | P3H2 | PAX6 | PEX11B | PEX16 | PEX5 |
| PEX7 | PITX3 | POLG | POLG2 | PXDN | RAB18 |
| RAB3GAP1 | RAB3GAP2 | RD3 | RDH11 | RDH12 | RECQL4 |
| RNASEH1 | RNLS | RPE65 | RPGRIP1 | RRM2B | SC5D |
| SEC23A | SIL1 | SLC16A12 | SLC25A4 | SLC2A1 | SLC33A1 |
| SORD | SPATA7 | SRD5A3 | SREBF2 | TBC1D20 | TDRD7 |
| TEAD1 | TMEM114 | TMEM70 | TULP1 | UNC45B | VIM |
| WFS1 | WFS1 | WRN |  |  |  |

57 genes associated with congenital cataract are marked with grey color.

Following sequencing, raw image files were processed using Bcl2Fastq software (Bcl2Fastq 2.18.0.12, Illumina, Inc.)for base calling and raw data generation.

Low-quality variations were filtered out using a quality score ≥20. Short Oligonucleotide Analysis Package (SOAP) aligner software (SOAP2.21; soap.genomics.org.cn/soapsnp.html) was then used to align the clean reads to the reference human genome. After PCR duplicates were removed, SNPs were identified by the GATK program (http://www.broadinstitute.org/gsa/wiki/index.php/Home_Page) and the SOAPsnp program (http://soap.genomics.org.cn/soapsnp.html). Identified SN-Ps and InDels were annotated using the Exome-assistant program (http://122.228.158.106/exomeassistant).

DNA samples from other individuals of the families were used to validate all mutations identified by using Sanger sequencing on ABI3730XL sequencing (Applied Biosystems Inc., CA). The coding regions of the candidate genes (MYH9, CRYBA4(c.169T>C), RPGRRIP1, WFS1, CRYBA4(c.26C>T), EPHA2, and PAX6) were amplified by polymerase chain reaction with the primers listed in the **Table 2** and screened by using bidirectional sequencing, which was finally analyzed with Chromas 2.33 and compared to the reference sequences in the NCBI database.

**Table 2.**
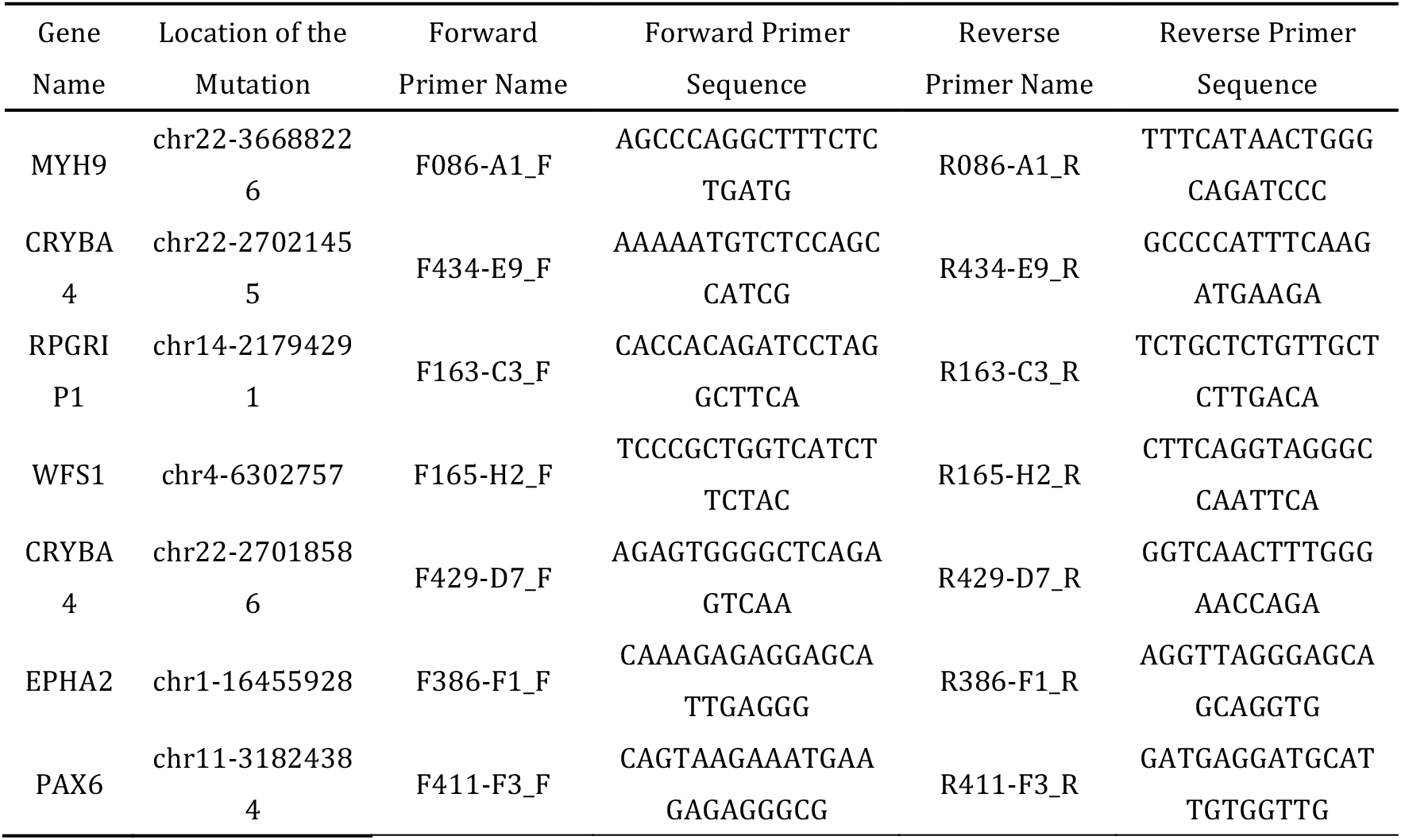
Primer Sequences for candidate genes (MYH9, CRYBA4(c.169T>C), RPGRRIP1, WFS1, CRYBA4(c.26C>T), EPHA2, and PAX6) amplification and sequencing

### Bioinformatics analysis

Based on the results we got from the mutation analysis, we conducted several bioinformatics analysis. The possible impact of an amino acid substitution on the structure and function of the protein was predicted by PROVEAN (Protein Variation Effect Analyzer, http://provean.jcvi.org/index.php) [11,12], Mutation Taster (http://www.mutationtaster.org) [13], Polymorphism and Phenotyping version 2 (http://genetics.bwh.harvard.edu/pph2/)[14] and GERP++ [15].

## Results

### Clinical findings

According to the family history and medical history, no individual had other systemic disease that could be related to cataract or other ophthalmic disease. In this study, we have identified 27 ADCC patients (25 alive) in six Chinese Families **(Figure 2**), including 9 males and 18 females (16 alive). Among them there are some patients whose opacified lens can be observed at a very young age (including patients in the Family 2, 4, 5and 6). The youngest patient we found in the six families was 4 months old. Her parents observed an opacified lens for the first time when she was only 2 months old. The non-transparent area deteriorated quickly during the past 2 months and eventually expanded to the whole lens (**Figure 2G**). However, patients in the Family 1 and 3 gradually showed the symptoms of ADCC after 11 years old.

It is worth noting that the proband of the Family 1 and her mother (suffering from ADCC) got a very different clinical feature that led to different influence on their visual acuity. The proband of the Family 1 is an 13-year-old girl with irregular nuclear cataract in both of her eyes (**Figure 2A**). Her visual acuity is 20/200 OU. However her mother with the same mutation in MYH9 gene develops a very mild symptom of ADCC. Her spot-like opacity is in the peripheral area of the lens (**Figure 2B**) which cause little damage to her visual acuity (20/25 OU).

**Table 3.**
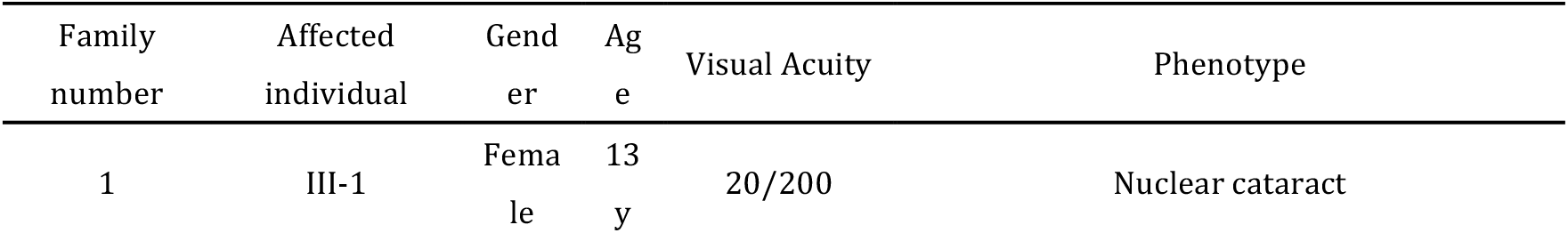
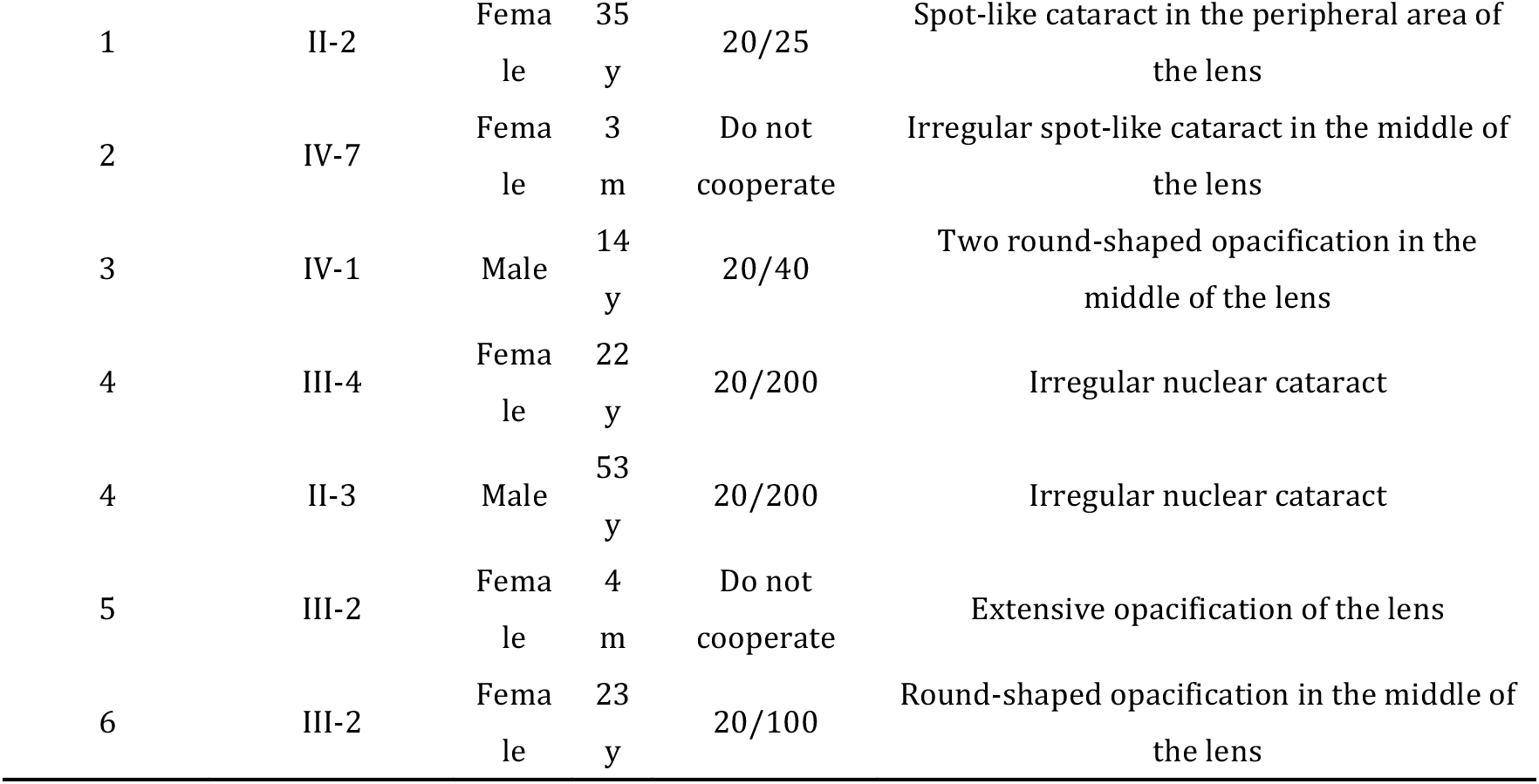
Clinical features of some affected individuals.

**Figure 2.**
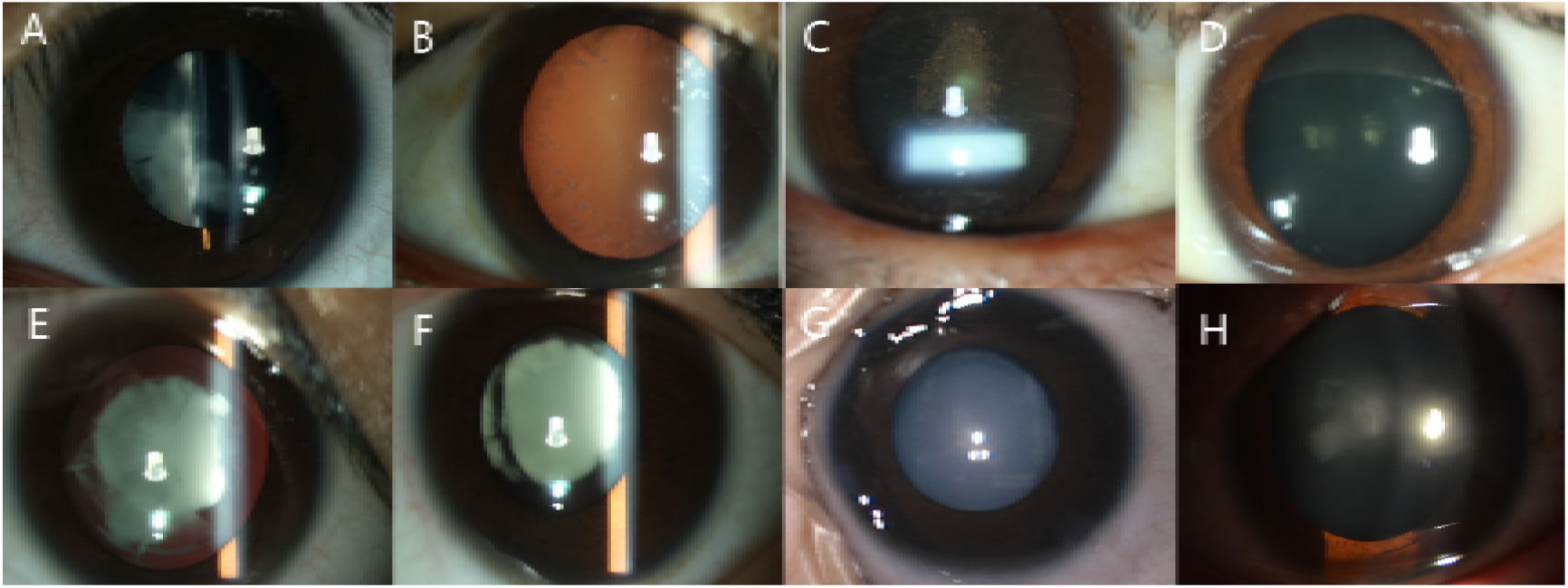
Slit lamp photograph of the family members: A. Proband of Family 1 (III-1); figure shows irregular opacities in the lens. B. II-2 in Family 1, the mother of the proband. C. Proband of Family 2 (IV-7). D. Proband of Family 3 (IV-1). E. Proband of Family-4 (III-4). F. II-3 in Family-4, the father of the proband. G. Proband of Family 5 (III-2) H. Proband of Family 6 (III-2)

### Mutation screening and Bioinformatics analysis

We conducted high throughput screening on the blood samples of all the probands. Compared with normal gene sequences, every proband had more than 3,000 nucleotide changes in hereditary eye disease related genes. Some of these nucleotide changes did not change amino acid sequence or accord with the hereditary model of ADCC and thus were excluded. It is worth noting that the genes associated with ADCC were screened out. Among them, the variant in the MYH9, RPGRRIP1, WFS1, EPHA2, and PAX6 of the probands were thought to be pathogenic possibly. After Sanger sequencing were conducted on the blood samples of other individuals in the families, it was found that the heterozygous MYH9 c.4150G>C, CRYBA4 c.169T>C, RPGRIP1 c.2669G>A, WFS1 c.1235T>C, CRYBA4 c.26C>T, EPHA2 c.2663+1G>A and PAX6 c.11–2A>G mutations were only detected in affected individuals, but didn’t present in unaffected individuals in the families. All these mutations were not detected in 100 controls and were considered to be related to ADCC.

#### A novel damaging missense mutation: MYH9 c.4150G>C mutation

The c.4150G>C mutation in MYH9 was identified in the Chinese Family 1 (**Figure 3A**). This missense mutation led to an amino acid change from Glutamate (E) to Glutamine (Q) at the codon 1384 (p.E1384Q) in the myosin heavy chain 9, also known as non-muscle myosin heavy chain IIa (NMMHC-IIA).

**Figure 3.**
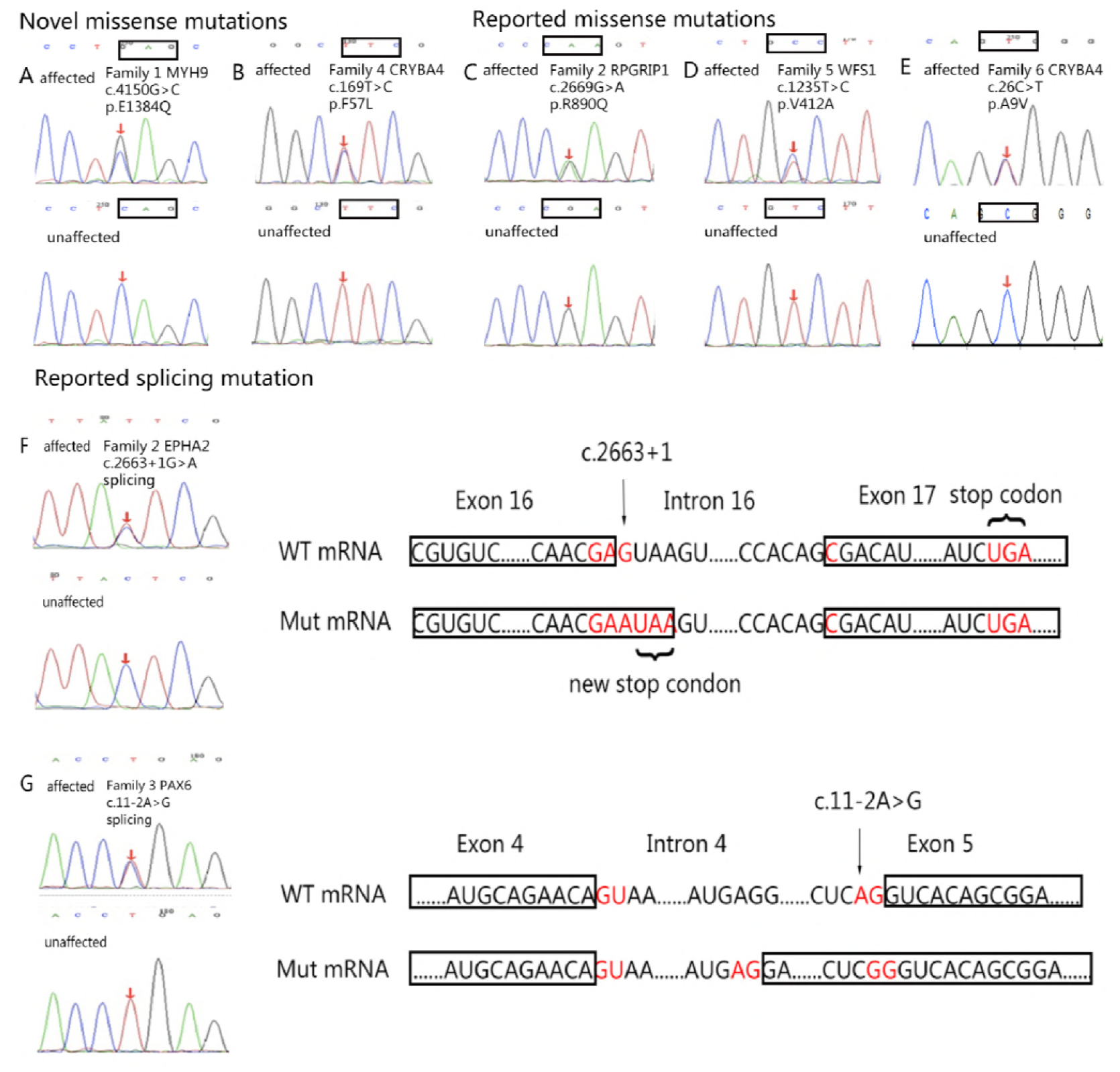
DNA sequences of affected and unaffected individuals (codons that are involved in the mutations are indicated with rectangles) A. MYH9 gene (c.4150G>C) B. CRYBA4 gene (c.169T>C) C. RPGRIP1 gene (c.2669G>A) D. WFS1 gene (c.1235T>C) E. CRYBA4 gene (c.26C>T) F. EPHA2 gene (c.2663+1G>A); the splicing mutation is in the first site of Intron 16, which leads to the change from Aspartic acid (D, encoded by GAC) to glutamate (E, encoded by GAA) as well as the creation of a new stop codon. This new stop codon causes the loss of 34 amino acids encoded by exon 17. G. PAX6 gene (c.11-2A>G); the c.11-2A>G mutation in PAX6 leads to the change from Serine (S, encoded by AGU) to Arginine (R, encoded by AGA) and an addition of 56 more amino acids.

According to the analysis of Mutation Taster, Glutamate (E) was highly conserved at the codon 1384 of MYH9 gene in different species (**Figure 4A**). Changing to Glutamine (Q) at this codon was predicted to be pathogenic by Mutation Taster (“Disease causing”), PROVEAN (“Deleterious” with the score of –2.64), SIFT (“Damaging” with the score of 0.023) and PolyPhen-2 (“Probably damaging” with the score of 1.000, sensitivity: 0.00; specificity: 1.00).

**Figure 4.**
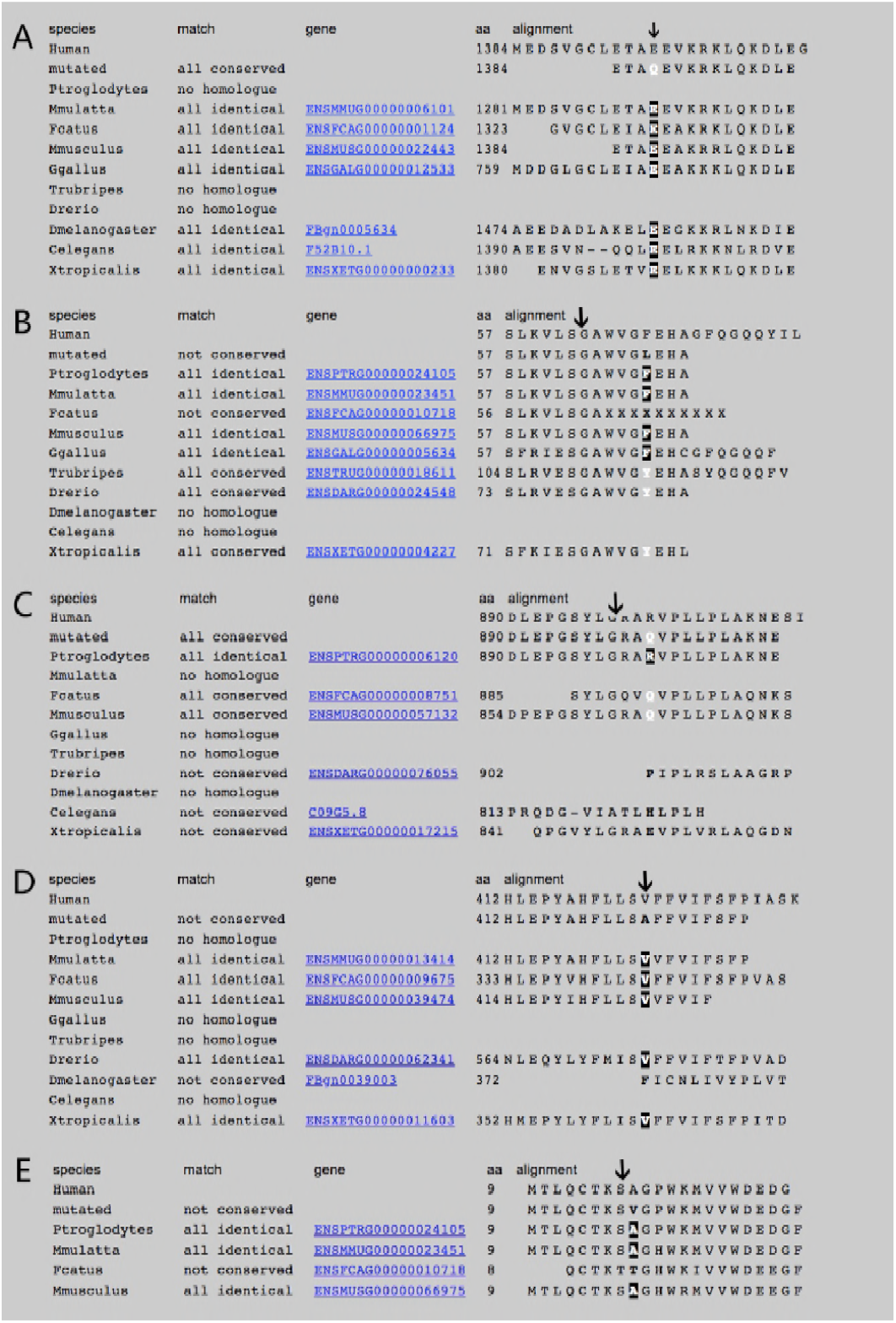
A. Glutamate (E) at the position 1384 of myosin heavy chain 9 is highly conserved in different species (indicted by an arrow). B. Phenylalanine (F) at the position 57 of crystalline beta A4 is conserved in different species (indicted by an arrow). C. Arginine (R) at the position890 of RPGRIP1 is not highly conserved in most species (indicted by an arrow). D. Valine (V) at the position 412 of WFS1 is conserved in different species (indicted by an arrow). E. Alanine (A) at the position 9 of crystalline beta A4 is conserved in different species (indicted by an arrow)

#### A novel damaging missense mutation: CRYBA4 c.169T>C mutation

The c.169T>C mutation in CRYBA4 was identified in the Chinese Family 4 **(Figure 3B**). This missense mutation led to an amino acid change from Phenylalanine (F) to Leucine (L) at the codon 57 (p.F57L) in the crystallin beta A4 (CRYBA4), a member of crystallin family.

According to the analysis of Mutation Taster, Phenylalanine (F) was conserved at the codon 57 of CRYBA4 gene in different species (**Figure 4B**). This mutation was then predicted to be pathogenic by Mutation Taster (“Disease causing”), PROVEAN (“Deleterious” with the score of –5.08), SIFT (“Damaging” with the score of 0.002) and PolyPhen-2 (“Probably damaging” with the score of 0.968, sensitivity: 0.74; specificity: 0.96). The 3D models of wild type and mutation type of CRYBA4-encoding protein were then made by Swiss model (**Figure 5**).

#### A novel neutral missense mutation: RPGRIP1 c.2669G>A mutation

The RPGRIP1 c.2669G>A in the Family 2 was identified in the Family 2 (**Figure 3C**

). This missense mutation can cause a “Neutral”(PROVEAN prediction with a score of 1.41) and “Tolerated”(SIFT prediction with a score of 0.867) amino acid change from Arginine (R) to Glutamine (Q) at the codon890 (p.R890Q) in the RPGR interacting protein 1. The amino acid is not highly conserved in different species (**Figure 4C**). Mutation Taster gave a prediction of “polymorphism” and PolyPhen-2 analysis scored this mutation with 0.000 (sensitivity: 1.00; specificity: 0.00).

#### A reported missense mutation: WFS1 c.1235T>C mutation

We identified the c.1235T>C mutation in WFS1 gene in the Chinese Family 5 **(Figure 3D**). This missense mutation could eventually cause an amino acid change from Valine (V) to Alanine (A) at the codon 412 (p.V412A).

The analyzing result of Mutation Taster showed that Valine (V) was conserved at the codon 412 of WFS1 in different species (**Figure 4D**) and the mutation was then predicted to be pathogenic by Mutation Taster (“Disease causing”), SIFT (“Damaging” with the score of 0.021) and PolyPhen-2 (“Probably damaging” with the score of 0.981, sensitivity: 0.75; specificity: 0.96). However, PROVEAN gave the prediction of “Neutral” with the score of –2.29.

This mutation was first reported by Choi et al.[16] as a candidate gene for familial nonsyndromic hearing loss. But according to what we found in the Chinese family 5, this mutation could also cause ADCC.

#### A reported missense mutation: CRYBA4 c.26C>T mutation

We identified the c.26C>T mutation in CRYBA4 gene in the Chinese Family 6 **(Figure 3E**), which could eventually cause an amino acid change from Alanine (A) to Valine (V) at the codon 9 (p.A9V). This mutation has been reported by Sun et al.[17] and Zhai et al.[18].

We all found clinical features of ADCC in these families. But the bioinformatics analysis results point out this mutation may not be pathogenic. According to the Mutation Taster, Alanine (A) was conserved in different species (**Figure 4E**). But the mutation was then predicted to be “polymorphism”. Besides, the mutation of the amino acid (p.V412A) was then predicted by PROVEAN, SIFT and PolyPhen-2.

The PROVEAN prediction result showed that this mutation was “Netural” with the score of –0.27. The SIFT predicted the mutation as a “Tolerated” mutation with the score of 0.502. PolyPhen-2 analysis gave a “Benign” prediction and scored this mutation with 0.009 (sensitivity: 0.96; specificity: 0.77).

Zhai et al.[18] assumed that this mutation was cosegregated with ADCC. We approve this hypothesis but more experiments should be conducted to verified it.

#### A reported splicing mutation: EPHA2 c.2663+1G>A mutation

After mutation screening, we identified a splice donor site mutation in EPHA2 (c.2663+1G>A, chr-16455928)(**Figure 3F and 3H**) in the Family 2. The result of the screening showed that the mutation is in the first site of Intron 16 and is predicted to change the Aspartic acid (D) to Glutamate (E). Also it is predicted to create a new stop codon which causes the loss of 34 amino acids encoded by exon 17[19].

Though we cannot predict the severe consequence of this mutation through SIFT, PROVEAN and PolyPhen2, we can still tell the damage of this mutation by using Mutation Taster and GERP++ prediction. The mutation in EPHA2 got a GERP score of 5.77, which is more than 3 and is considered to be a conservative site. The Mutation Taster gave a disease-causing prediction by pointing out that there were sequence motif loss and loss of protein feature.

We searched related papers for this mutation and found that it had been reported by Bu, et al [19]. The ID of mutation site in EPHA2 is chr1–16455928. But we found that the truly mutation site in EPHA2 was not c.2825+1G>A as the paper said but c.2663+1G>A as we detected since it fit the screening result and the sequence we searched from the NCBI.

#### A reported splicing mutation: PAX6 c.11–2A>G mutation

The mutation in PAX6 was also identified as a splicing mutation in the intron 4, which is located 2 sites before the beginning of exon 5. This mutation is predicted to change Serine (S, encoded by AGU) to Arginine (R, encoded by AGA). Also, it is predicted to cause the addition of 56 more amino acids (**Figure 3G and 3I**) [20].

SIFT, PROVEAN and PolyPhen2 cannot give the prediction of the mutation in PAX6. But according to the GERP++ score of 4.9 and Mutation Taster score of 1, we can tell that the site is highly conserved in different species. The Mutation Taster gives a “Disease causing” prediction. Here, we present an image of 3D models of PAX6 wild type and mutation type. It is obvious to find that the additional amino acids form an abnormal long “tail” which may influence the function of PAX6 (**Figure 5**). We searched related papers for this mutation and found that this mutation had been reported by Churchill, et al.[20].

**Figure 5.**
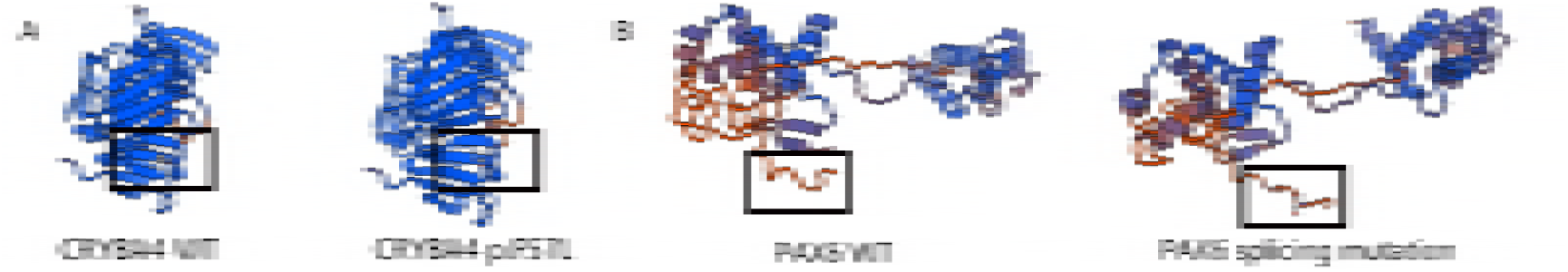
3D models of wild type and mutation type of CRYBA4-encoding protein and PAX6-encoding protein. The locations of the mutations are indicted by rectangles. A. CRYBA4 wild type and CRYBA4 p.F57L mutation B. PAX6 wild type and PAX6 splicing mutation

## Discussion

We used a specified panel that included 135 target cataract-associated genes to conduct screening tests on six Chinese families with congenital cataract. Our study detected seven suspected mutations in six genes, including two novel deleterious missense mutations (MYH9 c.4150G>C and CRYBA4 c.169T>C).

### A novel damaging mutation: MYH9 c.4150G>C mutation

The c.4150G>C variant is a missense mutation in the exon31 of MYH9 and leads to the substitution of Glutamate to Glutamine at the codon1384 (p.E1384Q) in the MYH9-encoded myosin heavy chain 9, also known as non-muscle myosin heavy chain IIa (NMMHC-IIA). Moreover, this mutation was not detected in 100 normal controls suggesting that it is a pathogenic mutation rather than a polymorphism. All the bioinformatics analysis suggested that this mutation was a disease causing mutation. Thus, these results support that the congenital cataract in the Family 1 is caused by the mutation in the exon31 of MYH9 gene.

Mutations on MYH9 may cause MYH9 disorder, which is characterized by congenital macrothrombocytopenia complicated with young-adult-onset sensorineural hearing loss, nephropathy and congenital cataract. However, the etiologies of the diseases above are still unknown [21]. So far, there have been several reports of mutations in MYH9 gene (**Table 3**). According to Pecci A et al. cataract did not commonly occur in family with MYH9 disorder and were only observed in 43 of 235 (18%) patients. Among them, there were only 4 congenital forms [22]. In our research, the Family 1 that we recruited had the ocular manifestation of congenital cataract without renal complication and hearing loss, whose symptoms were extremely mild.

**Table 3.**
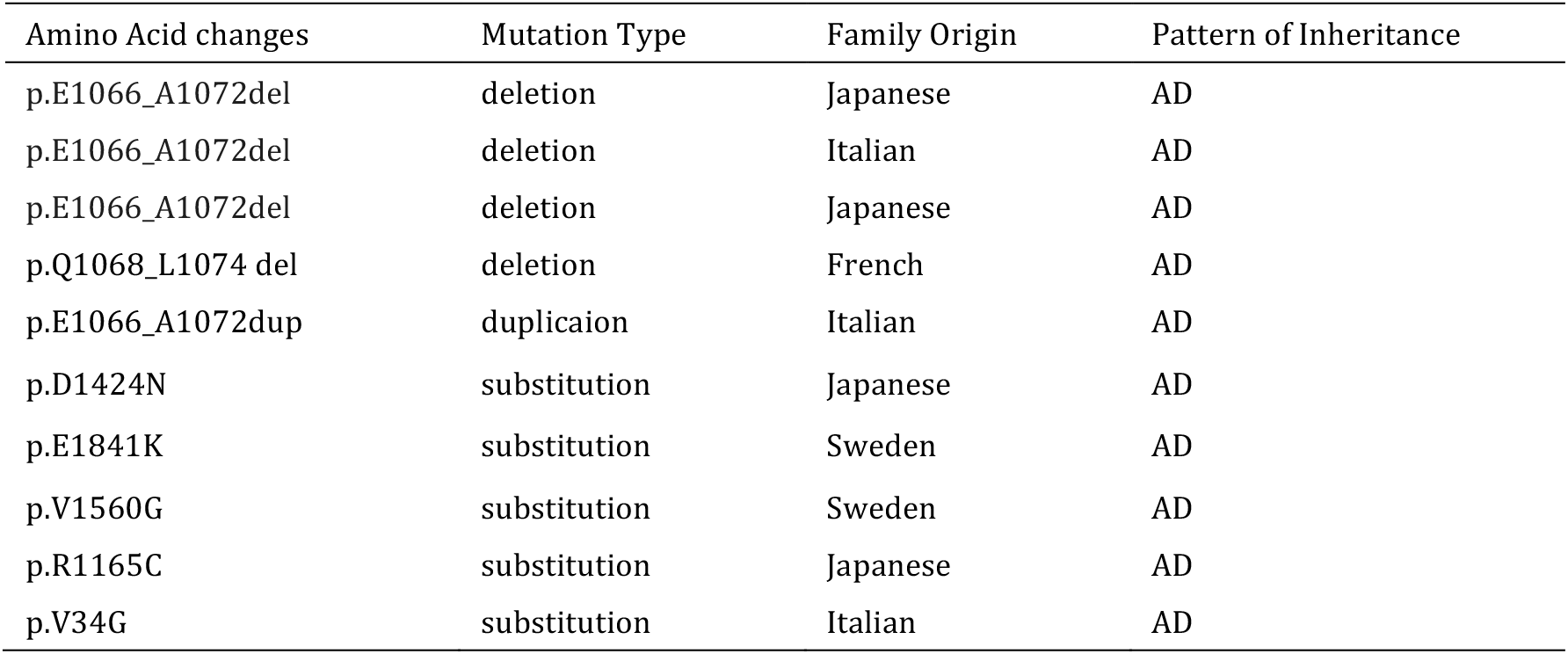
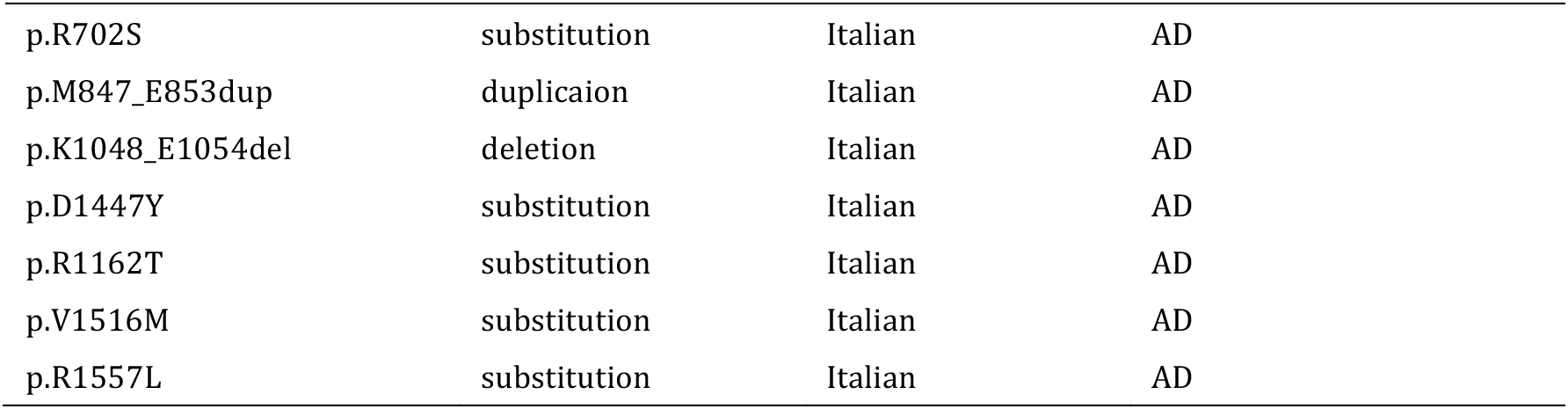
Mutation in MYH9 associated with congenital cataract

The myosin heavy chain 9 gene, mapped on chromosome 22q12.3, includes 47 exons, which encode non-muscle myosin heavy chain IIA. The transcription of this gene starts with the first ATG of the open reading frame in exon 2 and stops at the stop codon in exon 41. The mutation we identified was in the exon 31, which probably affected the tail domain of the NMMHC-IIa. According to Alessandro Pecci et al., there is a strong correlation between genotype and clinical aspects in MYH9 disorder [23]. Moreover, mutations in the rod-tail domain are associated with a mild phenotype [24]. Thus, we predict that mutations in the exon 31 will lead to hereditary disease, but in an extremely mild phenotype, which is proved by the manifestation of the Family 1 we recruited. Moreover, this mutation was not detected in 100 normal controls suggesting that it is a pathogenic mutation rather than a polymorphism. All the bioinformatics analysis suggested that this mutation was a disease causing mutation. Thus, these results support that the congenital cataract in the Family 1 is caused by the mutation in the exon31 of MYH9 gene.

### A novel damaging mutation: CRYBA4 c.169T>C mutation

The missense mutation (c.169 T>C) in CRYBA4 leads to the amino acid change from Phenylalanine (F) to Leucine (L) at the codon 57 (p.F57L) in the crystallin beta A4 (CRYBA4), a member of crystallin family. This mutation was not detected in 100 normal controls and was predicted to be pathogenic by different bioinformatics analysis.

Crystallin beta A4 (CRYBA4) belongs to beta-crystallin family. The beta-crystallin family(35%), alpha-crystallin (40%) and gamma-crystallin are six major members of crystallin family found in mammalian lens[25]. Crystallins are made very stable in lens to keep transparency and refractive ability. The beta-crystallin includes 7 protein forms, each of which is encoded by six genes (CRYBA1, CRYBA2 and CRYBA4; CRYBB1, CRYBB2 and CRYBB3). Among them, CRYBA4 encodes 196 amino acids and pathogenic mutations are usually in the exon 2 and exon 4 of this gene (**Table 4**). According to Bilingsley et al.[25]and Zhou [26], the mutation in exon 4 play an important role in the onset of congenital cataract. The novel mutation we detected is also in this area.

**Table 4.** Mutation in CRYBA4 associated with congenital cataract

| Amino Acid changes | Mutation Type | Family Origin | Pattern of Inheritance |
| --- | --- | --- | --- |
| p.L69P | substitution | India | AD |
| p.F94S | substitution | India | AD |
| p.G2D | substitution | China | AD |
| p.D3Y | substitution | Honduras | AD |
| p.L11S | substitution | Denmark | AD |
| p.T19M | substitution | India | AD |
| p.V28M | substitution | India | AD |
| p.F32L | substitution | China | AD |
| p.R33L | substitution | India | AD |
| p.V44M | substitution | China | AD |
| p.V44M | substitution | USA | AD |
| p.W45S | substitution | China | AD |
| p.D47N | substitution | China | AD |
| p.P59L | substitution | USA | AD |
| p.A9V | substitution | China | AD |
| p.G147V | substitution | Pakistani | AD |

As is illustrated in the picture of 3D models of CRYBA4 wild type and mutation type (**Figure 5**), Phenylalanine (F) at the codon 57 is replaced by Leucine (L). The mutation is located in the beta-sheet. Beta-sheet is stabilized by hydrogen bond. The substitution from Phenylalanine (F) to Leucine (L) may break the hydrogen bond, resulting in the damage of the stabilization of the secondary structure of the CRYBA4-encoding protein.

Considering the differences between splicing mutation and mutation that change amino acids and structure of proteins, we conducted different bioinformatics analysis to figure out the consequences of these mutations. We find that comparing to the amino acid changing mutations, splicing mutations may be more dangerous because they may change the transcription and translation by creating new stop codons or adding new amino acid sequences. These changes can cause loss of motif or change of structure and influence the function of proteins eventually.

Through the study of these families, we should confirm that the detecting strategies we used to reveal a novel mutation is notable. Considering the number of individuals in the families and the number of cataract-associated genes we should detect, using targeted exome sequencing to identify the mutation of the proband and Sanger sequencing for the rest are much more efficient and economical. Fortunately, we are cooperating with Reproductive Medicine Center of Peking University Third Hospital to see if we can conduct prenatal diagnosis to block the pathogenic mutation.

More information should be collect to verify whether the cataracts caused by these mutations are progressive. However, some details of the phenotype of the affected individuals of the families cannot be observed, as some cataracts have been removed by surgeries.

## Acknowledgements

We would like to thank our patients and their family members for their participation. This study was funded by the National Science and Technology Major Project (2018ZX10101004).

## Reference

1. Hejtmancik J. F., 2008 Congenital cataracts and their molecular genetics. Semin Cell Dev Biol 19(2):134–149.

2. Robinson G. C., Jan J. E., Kinnis C., 1987 Congenital ocular blindness in children, 1945 to 1984. Am J Dis Child 141(12):1321–1324.

3. Shiels A. and J. F. Hejtmancik., 2007 Genetic origins of cataract. Arch Ophthalmol 125(2):165–173.

4. Huang B. and W. He., 2010 Molecular characteristics of inherited congenital cataracts. Eur J Med Genet 53(6):347–357.

5. Francis P. J., Berry V., Bhattacharya S. S., Moore A. T., 2000 The genetics of childhood cataract. J Med Genet 37(7):481–488.

6. Apple D. J., Ram J., Foster A., Peng Q., 2000 Elimination of cataract blindness: a global perspective entering the new millenium. Surv Ophthalmol 45 Suppl 1:S1–196.

7. Santana A. and M. Waiswo., 2011 The genetic and molecular basis of congenital cataract. Arq Bras Oftalmol 74(2):136–142.

8. Gill D., Klose R., Munier F. L., McFadden M., Priston M., et al., 2000 Genetic heterogeneity of the Coppock-like cataract: a mutation in CRYBB2 on chromosome 22q11.2. Invest Ophthalmol Vis Sci 41(1):159–165.

9. Liu Q., Wang K. J., Zhu S. Q. 2014 A novel p.G112E mutation in BFSP2 associated with autosomal dominant pulverulent cataract with sutural opacities. Curr Eye Res 39(10):1013–1019.

10. Kunishima S., Kojima T., Matsushita T., Tanaka T., Tsurusawa M., et al., 2001 Mutations in the NMMHC-A gene cause autosomal dominant macrothrombocytopenia with leukocyte inclusions (May-Hegglin anomaly/Sebastian syndrome). Blood 97(4):1147–1149.

11. Choi Y. and A. P. Chan. 2015 PROVEAN web server: a tool to predict the functional effect of amino acid substitutions and indels. Bioinformatics 31(16):2745–2747.

12. Choi Y., Sims G. E., Murphy S., Miller J. R., Chan A. P., 2012 Predicting the functional effect of amino acid substitutions and indels. PLoS One 7(10):e46688.

13. Schwarz J. M., Rodelsperger C., Schuelke M., Seelow D., 2010 MutationTaster evaluates disease-causing potential of sequence alterations. Nat Methods 7(8):575–576.

14. Adzhubei I. A., Schmidt S., Peshkin L., Ramensky V. E., Gerasimova A., et al., 2010 A method and server for predicting damaging missense mutations. Nat Methods 7(4):248–249.

15. Davydov E. V., Goode D.L., Sirota M., Cooper G.M., Sidow A., et al., 2010 Identifying a high fraction of the human genome to be under selective constraint using GERP++. PLoS Comput Biol 6(12): e1001025.

16. Choi B. Y., Park G., Gim J., Kim A. R., Kim B. J., et al., 2013 Diagnostic application of targeted resequencing for familial nonsyndromic hearing loss. PLoS One 8(8): e68692.

17. Sun W., Xiao X., Li S., Guo X., Zhang Q., 2014 Exome sequencing of 18 Chinese families with congenital cataracts: a new sight of the NHS gene. PLoS One 9(6): e100455.

18. Zhai Y., Li J., Yu W., Zhu S., Yu Y., et al., 2017 Targeted Exome Sequencing of Congenital Cataracts Related Genes: Broadening the Mutation Spectrum and Genotype-Phenotype Correlations in 27 Chinese Han Families. Sci Rep 7(1): 1219.

19. Bu J., He S., Wang L., Li J., Liu J., Zhang X., 2016 A novel splice donor site mutation in EPHA2 caused congenital cataract in a Chinese family. Indian J Ophthalmol 64(5): 364–368.

20. Churchill A. J., Hanson I. M., Markham A. F., 2000 Prenatal diagnosis of aniridia. Ophthalmology 107(6): 1153–1156.

21. Aoki T., Kunishima S., Yamashita Y., Minamitani K., Ota S., 2018 Macrothrombocytopenia With Congenital Bilateral Cataracts: A Phenotype of MYH9 Disorder With Exon 24 Indel Mutations. J Pediatr Hematol Oncol 40(1):76–78.

22. Pecci A., Klersy C., Greseie P., Lee K. J., De Rocco D., et al., 2014 MYH9-related disease: a novel prognostic model to predict the clinical evolution of the disease based on genotype-phenotype correlations. Hum Mutat 35(2):236–247.

23. Pecci A., Panza E., Pujol-Moix N., Klersy C., Di Bari F., et al., 2008 Position of nonmuscle myosin heavy chain IIA (NMMHC-IIA) mutations predicts the natural history of MYH9-related disease. Hum Mutat 29(3):409–417

24. Pecci A., Panza E., De Rocco D., Pujol-Moix N., Girotto G., et al., 2010 MYH9 related disease: four novel mutations of the tail domain of myosin-9 correlating with a mild clinical phenotype. Eur J Haematol 84(4):291–297.

25. Billingsley G., Santhiya S. T., Paterson A. D., Ogata K., Wodak S., et al., 2006 CRYBA4, a novel human cataract gene, is also involved in microphthalmia. Am J Hum Genet 79(4): 702–709.

26. Zhou G., Zhou N., Hu S., Zhao L., Zhang C., et al., 2010 A missense mutation in CRYBA4 associated with congenital cataract and microcornea. Mol Vis 16: 1019–1024.

